# Functional Sensitivity and Mutational Robustness of Proteins

**DOI:** 10.1101/2020.06.12.148304

**Authors:** Qian-Yuan Tang, Testsuhiro S. Hatakeyama, Kunihiko Kaneko

## Abstract

Sensitivity and robustness appear to be contrasting concepts. However, natural proteins are robust enough to tolerate random mutations, meanwhile be susceptible enough to sense environmental signals, exhibiting both high functional sensitivity (i.e., plasticity) and mutational robustness. Uncovering how these two aspects are compatible is a fundamental question in the protein dynamics and genotype-phenotype relation. In this work, a general framework is established to analyze the dynamics of protein systems under both external and internal perturbations. We introduce fluctuation entropy for the functional sensitivity and the spectrum entropy for the mutational robustness. The compatibility of sensitivity and robustness is analyzed by the optimization of two entropies, which leads to the power-law vibration spectrum of proteins. These power-law behaviors are confirmed extensively by protein data, as a hallmark of criticality. Moreover, the dependence of functional sensitivity and mutational robustness on the protein size suggests a general evolutionary constraint for proteins with different chain lengths. This framework can also establish a general link of the criticality with robustness-plasticity compatibility, both of which are ubiquitous features in biological systems.

## I. INTRODUCTION

Proteins are highly dynamic molecules in living cells [1–4]. As a living system, a protein molecule is subjected to external and internal perturbations. For proteins in the solutions, small external perturbations from the milieu can be felt consistently by every other residue within the entire protein. Even weak and transient perturbations can trigger large conformational changes [5–8]. In this way, proteins can sense external signals ranging from metal ions to other biomolecules, which are functionally beneficial.

Meanwhile, internal perturbations also affect proteins. Numerous mutations occur intermittently in gene sequences encoding amino acid residues of proteins. In the “sequence-structure-dynamics-function” paradigm [9, 10], mutations may change the native protein structure; consequently, the equilibrium dynamics and corresponding biological functions will be affected. However, accumulating evidence has shown that protein molecules can tolerate numerous types of amino acid substitutions and maintain their thermodynamic stability and folding pathways [11–15]. Besides, the principal components of protein equilibrium dynamics are highly robust to minor structural differences or detailed atomic-level interactions [9, 16–19]. These results demonstrate that the dynamics of proteins are highly robust to mutations.

Indeed, both high sensitivity to external perturbations and high robustness to internal perturbations are essential for protein systems [20]. Proteins without a high functional sensitivity cannot respond transiently or undergo large-scale conformational changes. Meanwhile, without high mutational robustness, the functional proteins with high fitness cannot be stably replicated or inherited. However, the former and latter concern changeability and non-changeability, respectively, and may direct us to different ends for systems with a single degree of freedom [21–24]. Noting that the proteins involve many degrees of freedom, the utilization of the multiple degrees of freedom raises the possibility to achieve both the two aspects. To resolve such a problem, the native-state dynamics and corresponding responses to noises and mutations may provide new perspectives on the genotype-phenotype relation of the proteins.

In this paper, with a simplified protein model, we establish a general framework for analyzing protein dynamics, which is subjected to external and internal perturbations. We introduce the fluctuation entropy and spectrum entropy to describe the functional sensitivity and mutational robustness, respectively. Then, from the perspective of the vibration spectrum, the interplay between functional sensitivity and mutational robustness of proteins is analyzed. In systems with low degrees of freedom, the two entropies act as consistent descriptors. However, in systems with high degrees of freedom, the interplay between functional sensitivity and mutational robustness will lead to a power-law eigenvalue distribution in the relaxation dynamics. Such a power-law distribution is supported by the vibration spectra of natural proteins. The corresponding power-law coefficient is closely associated with the packing density of protein molecules. The size dependence of functional sensitivity and mutational robustness suggests a general evolutionary constraint for proteins with different sizes.

## II. FUNCTIONAL SENSITIVITY AND MUTATIONAL ROBUSTNESS

### A. Model

Globular proteins fold into their native structure to perform their biological functions. For proteins fluctuating around their equilibrium configuration, the relaxations and fluctuations can be captured by linear models. By approximating the energy landscape around the native state in the quadratic form, potential energy

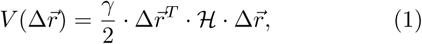

in which 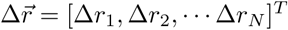 is a vector describing the magnitude of displacement of all *N* residues deviating from the native structure, *γ* is the spring constant, and ℋ is the second-order derivative of potential energy, known as the Hessian matrix. In this way, a protein molecule can be modeled into an elastic network, where amino acid residues (represented by nodes positioned at their C_*α*_ atom) are connected via elastic springs [25]. As a simple yet powerful model, elastic networks are applied widely for elucidating the functional dynamics [26, 27] and evolutionary constraints [28–31] of proteins.

Additionally, after assuming that all residue fluctuations are Gaussian variables distributed around their equilibrium coordinates, Gaussian Network Model (GNM) is introduced to describe the fluctuations around native protein structures [32, 33]. In the GNM, when the interaction strength between residues are assigned to be 0 or 1, the Hessian matrix ℋ can be reduced as the graph Laplacian, describing the topology of the network (see Appendix A). To have a better prediction of protein dynamics, one can also assign different weights for different residue pairs (see SM, Sec. 1.1). The structural fluctuations around the native state can then be described by a multivariate Gaussian distribution [17], given by

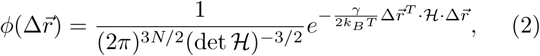

where *k*_*B*_ and *T* denote the Boltzmann constant and temperature, respectively.

For globular proteins, based on the GNM, the correlated motions of the proteins can be predicted (see Appendix B). By conducting normal mode analysis (NMA), one can obtain essential information about thermal fluctuations, large-scale conformational dynamics, and linear responses of native proteins [34–37]. Upon the eigen-value decomposition of the Hessian matrix ℋ, the normal modes described by eigenvectors 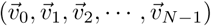 and corresponding eigenvalues (0 = *λ*_0_ *< λ*_1_ ≤ *λ*_2_ ≤ …*λ*_*N*−1_) can be obtained. Notably, the matrix ℋ is positive semi-definite; there is always *λ*_0_ = 0, showing the translational invariance of the system. The native dynamics of proteins can then be described as a linear combination of independent normal modes. It is worth noting that, even for the large-scale nonlinear conformational changes, the normal modes given by elastic network models could have a high overlap with such deformations [27, 28].

### B. Functional sensitivity

Here, we introduce “functional sensitivity” to quantify that how much will an external perturbation leads to the deformations of a protein molecule. For native proteins, high functional sensitivity usually correlates with low structural robustness, and it is closely associated with the geometry at the native basin of the energy landscape. In NMA, the eigenvalue magnitudes provide information about the local curvature of the energy landscape along the direction of the corresponding eigenvectors [38]. An illustrative energy landscape is shown in Fig. 1A. Here, *λ*_*j*_ *< λ*_*k*_ indicates that the motion along 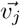 is more easily accessible, and lesser energy is needed to excite such deformations, as compared to that for the motion along the direction of 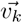. To quantify the functional sensitivity of a protein molecule, one shall measure the total “volume” of the accessible configuration space at the given perturbation level (temperature). To calculate that, we introduce the fluctuation entropy *S*_*F*_ of the protein molecule as the Shannon entropy (or differential entropy) [39–41] of the fluctuation distribution 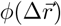.

**FIG. 1.**
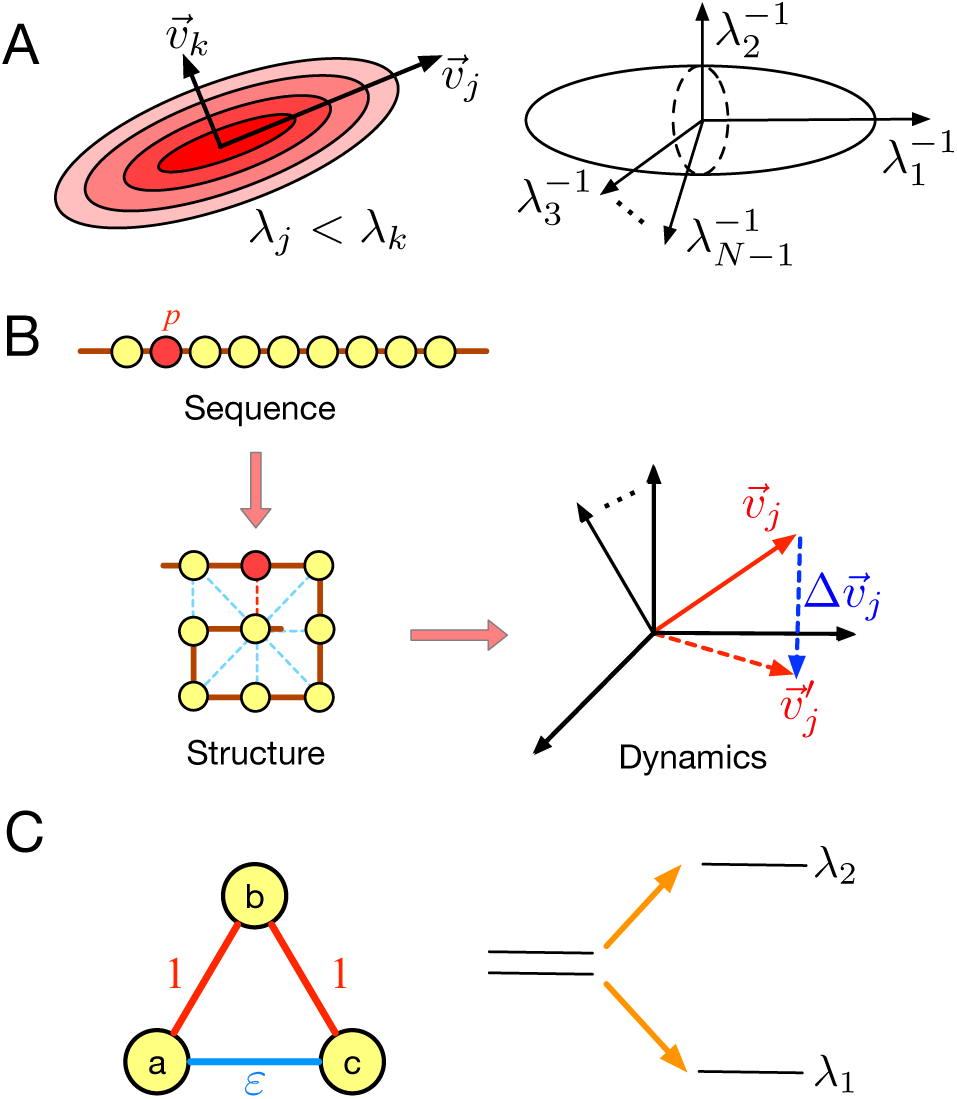
Illustration of functional sensitivity and mutational robustness. (A) Left: When *λ*_*j*_ *< λ*_*k*_, as compared to 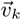, the motions along 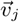 will be easier to access, and less energy is required for deformation. Right: The configuration space of the protein molecule around the native state is shown to be a high-dimension ellipsoid, in which the length of the *j*-th semi-axes is 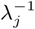. The fluctuation entropy is defined as the logarithm of this volume. (B) The original amino acid sequence folds into a specific native protein structure and exhibits specific functional dynamics. After the mutation on-site *p*, interactions between residue *p* and other neighboring residues change, and the dynamics of such a protein also change slightly. For example, the *j*-th eigenvector 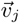 will transform into 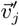 after such a mutation. The difference in such an eigenvector is 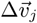. (C) An illustrative model of the three-bead chain is shown. Here, maximizing the eigengap can contribute to both functional sensitivity and mutational robustness.

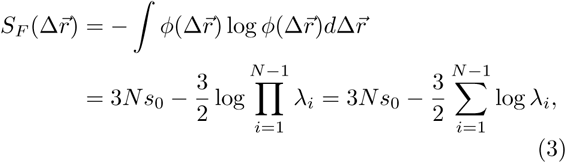

in which 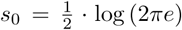. The derivation of Eq. (3) is listed in Sec. 1.2 of Supplementary Materials (SM). According to such an expression, the fluctuation entropy of protein molecules can be decomposed into two parts: a constant term and another term equal to the logarithm of the product 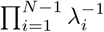. This product is also known as the pseudo-determinant of the matrix ℋ [42]. For the *j*-th normal mode, the 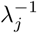 describes variance along the direction of the eigenvector 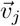. As illustrated in Fig. 1A, the pseudo-determinant can be understood as the volume of the configuration space; the fluctuation entropy is the logarithm of such a volume.

The physical meaning of the fluctuation entropy *S*_*F*_ is clear: when there are more eigenvalues with small magnitude, the corresponding slow-mode motions will be easier to activate. The fluctuation entropy of such a molecule is high, indicating that a small perturbation can lead to large fluctuations in such a system. It is worth noting that our definition of fluctuation entropy is consistent with other measures for system sensitivity, such as the condition number. Such consistency is discussed in the SM (Sec. 2). Besides, previous experimental studies also showed that the conformational entropy can contribute significantly to functionality of the proteins [43].

### C. Mutational robustness

In living cells, mutations happen from time to time. For a protein system, the mutations can be understood as internal perturbations. Mutations in the sequence may cause changes in protein structures. However, structural differences or residue-residue interactions do not necessarily imply that the dynamics of a protein are different [16–19]. For example, the structure of the apo and holo states of an allosteric protein may differ considerably, but a conformational transition can easily connect the two states. Besides, proteins as a product of evolution can tolerate numerous kinds of amino acid substitutions and maintain their native-state structures [11–13]. To quantify the consequential effects (changes in the phenotype) of mutations, in this work, we mainly focus on the robustness of the native dynamics of the protein [44–46].

For simplification, we take the dynamics of a protein as a linear combination of eigenvectors of the Hessian matrix ℋ. Recently, the quantitative assessment of the conservation of these eigenvectors has become a new topic in the study of protein evolution [47]. Here, we quantify the mutation-induced changes in protein native dynamics as the differences in eigenvectors (or the subspaces spanned by the eigenvectors) of such a system. As illustrated in Fig. 1B, a residue substitution at site *p* may change the interactions between it and its spatial neighbors. These changes will affect the elements of the Hessian matrix ℋ (see SM, Sec. 3.1). After the mutation, the Hessian matrix changes from ℋ to ℋ′, where ℋ′ = ℋ + *ϵM*. Here, the parameter *ϵ* denotes the perturbation magnitude, and *M* is a symmetric perturbation matrix that describes changes in inter-residue interactions. After the mutation, the eigenvalues of the Hessian matrix shift from *λ*_1_, *λ*_2_, … *λ*_*N*−1_ to 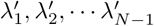, respectively, and the corresponding eigenvectors change from 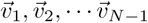 to 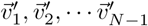, respectively.

To characterize the robustness of equilibrium dynamics in a protein molecule, one needs to check if the mutation can easily modify the eigenvectors (or the subspaces spanned by eigenvectors) which are closely associated with the functional motion of such a protein. If a protein can maintain its function-related eigenvectors or subspaces after a mutation, such a protein shows high mutational robustness in the evolutionary process.

Here, let us first consider the case that the functional motions can be described by a single eigenvector. Assume that the *j*-th eigenvector is associated with the functional motion of the protein, and *λ*_*j*−1_ *< λ*_*j*_ *< λ*_*j*+1_. According to the first-order perturbation theory [48], the perturbed eigenvector 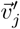 can be expressed as:

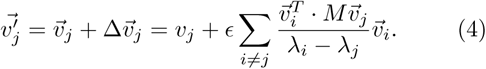

Therefore, for eigenvector 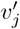, the square deviation is expressed as:

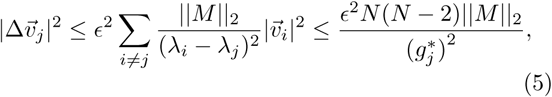

where ‖*M*‖_2_ denotes the *L*_2_-norm of matrix *M*, which is bound by the number of mutated contacts, and 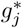 denotes the minimum gap between *λ*_*j*_ and its neighboring eigenvalues, say, 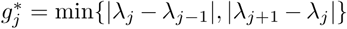. The magnitude of 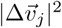 can quantify the robustness of vector 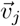 after perturbation. Larger 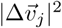 indicates more significant changes in the dynamics of the *j*-th eigenvector, which implies lower mutational robustness in protein dynamics. According to Eq.5, the upper bound of 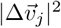 of the *j*-th eigenmode is determined by 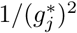; thus, a larger eigengap 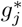 corresponds to a higher mutational robustness of the eigenvector 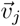.

In more complicated cases, the functional dynamics of a protein cannot be described by a single eigenvector, but needs to be expressed as a linear combination of a set of eigenvectors. Then, the above discussions shall be generalized for the mutational robustness of a linear subspace. Here, as an example, let us consider the linear subspace 𝒱_*j*_, spanned by the first *j* slow-mode eigenvectors 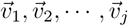. It can be concluded that the mutational robustness of the linear space 𝒱_*j*_ will be upper bounded by the eigengap |*λ*_*j*+1_ − *λ*_*j*_| (details in SM, Sec. 3.4).

### D. A system with low degrees of freedom

In previous discussions, we demonstrated that functional sensitivity and mutational robustness are closely related to the spectral properties of the elastic network. To illustrate the possible relationships between functional sensitivity and mutational robustness, let us first focus on a simple three-bead system with two degrees of freedom. As shown in Fig. 1C, the strength of the bonded interactions is 1, and the strength of the non-bonded interaction is *ε* (0 ≤ *ε* ≤ 1). By NMA, the non-zero eigen-values are found to be: *λ*_1_ = 1 + 2*ε* and *λ*_2_ = 3. To maximize the functional sensitivity, the non-bonded interaction *ε* shall be weakened, to ensure that the fluctuation entropy *S*_*F*_ ∼ −log(*λ*_1_*λ*_2_) is maximized. To maximize the mutational robustness of the first and second eigenvectors, the eigengap |*λ*_2_ − *λ*_1_| shall be maximized. When *ε* = 0, both functional sensitivity and mutational robustness reach maximum. This model illustrates that in systems with low degrees of freedom, maximizing the gaps between the two non-zero eigenvalues can contribute to both functional sensitivity and mutational robustness. For example, brain networks can be coarse-grained as block models, which only have low degrees of freedom. Previous studies have proved that larger gaps between eigenvalues, which correspond to the modular organization in the brain, may confer increased robustness to network perturbations and higher flexibility in learning [49].

## III. THE INTERPLAY BETWEEN FUNCTIONAL SENSITIVITY AND MUTATIONAL ROBUSTNESS FOR A SYSTEM WITH MANY DEGREES OF FREEDOM

For systems with high degrees of freedom, the interplay between functional sensitivity and mutational robustness is complicated. To maximize the functional sensitivity (fluctuation entropy *S*_*F*_) of a system, as illustrated in Fig. 2A, one will expect an eigenspectrum with *N* − 2 vibration modes, where eigenvalues are close to zero (*λ*_1_ ≈ *λ*_2_ ≈ … ≈ *λ*_*N*−2_ ≈ 0), and the largest eigenvalue *λ*_*N*−1_ is much larger than zero. In such a situation, the fluctuation entropy reaches the maximum, which ensures that external perturbations can easily trigger the motions of the molecule along the directions of the first *N* − 2 eigenvectors. However, the gaps between low-frequency modes are expected to be very small; thus, these low-frequency modes are not stable under mutations.

**FIG. 2.**
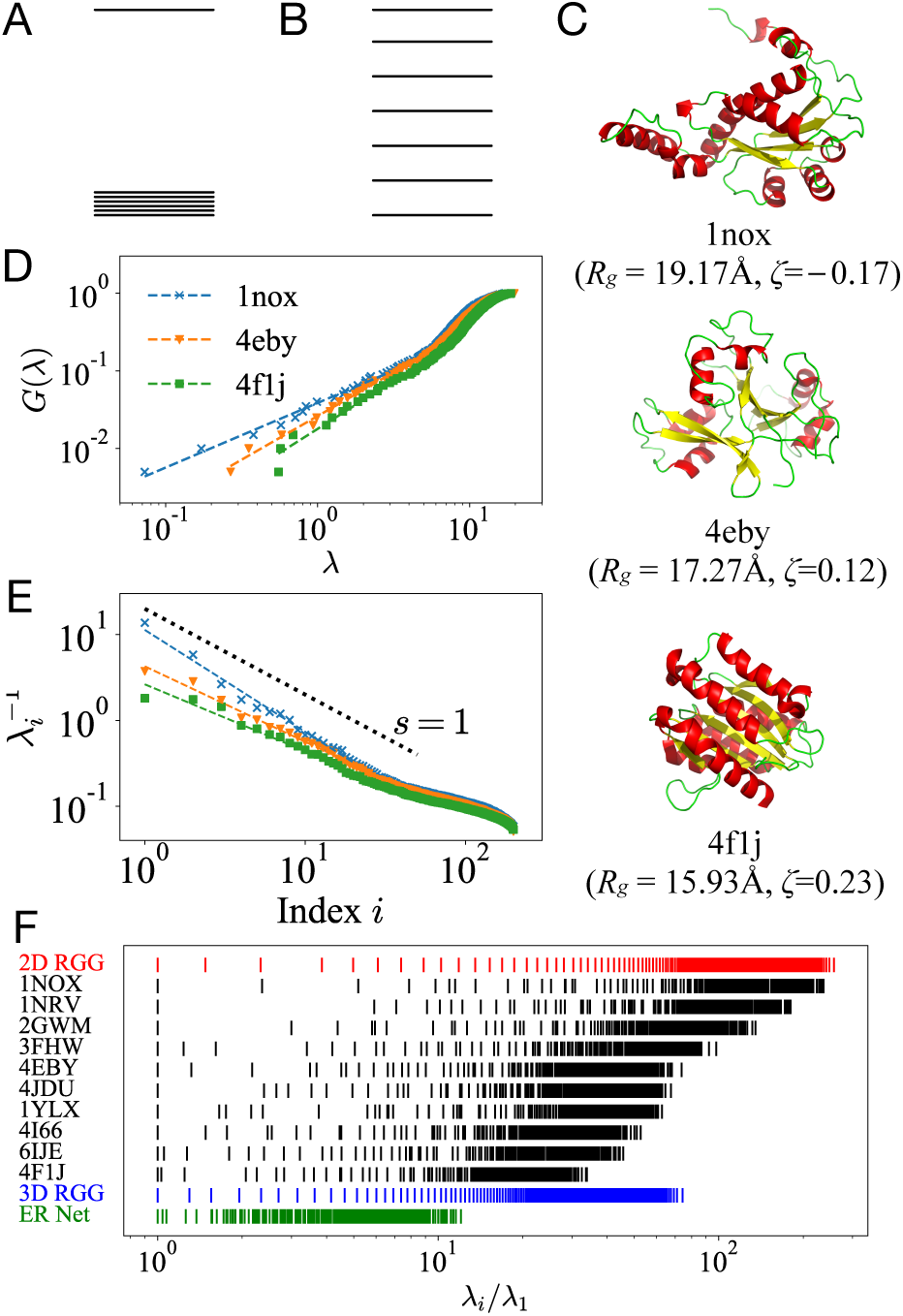
(A) The eigenvalue distribution that maximizes functional sensitivity. (B) The eigenvalue distribution that maximizes the mutational robustness of the linear space spanned by all the eigenvectors. (C) The cartoon structure of the three proteins (PDB code: 1NOX, 4EBY, and 4F1J) with the same chain length (*N* = 200). (D) The integrated eigen-value distribution function *G*(*λ*) and (E) the magnitude of the *i*-th mode 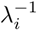 vs the index (rank) *i* for the three proteins is shown. The dashed lines in (D) and (E) show the fitting results based on the first 20 eigenvalues of the proteins. (F) The eigenvalue spectra of proteins with chain length *N* = 200 is shown. All eigenvalues are normalized to the lowest non-zero eigenvalue *λ*_1_, and the log-scale that clearly presents the differences between neighboring eigenvalues is selected. The spectra of proteins are compared with Erdős-Rényi networks and random geometric graphs (2D or 3D).

Let us consider the mutational robustness of the linear space 𝒱 spanned by all the eigenvectors 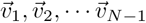. The mutational robustness of 𝒱 will be upper bounded by 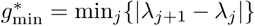 (proof in SM, Sec. 3.5). To maximize the robustness of eigenspace 𝒱 under mutations, the minimum eigengap 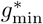 of the entire spectrum shall be maximized. This will lead to a uniform distribution of all the eigenvalues as long as the eigenvalues are bounded between *λ*_min_ and *λ*_max_. To quantify the uniformity of the eigenvalue distribution, we define spectrum entropy 𝒮 as the Shannon entropy of the eigenvalue distribution:

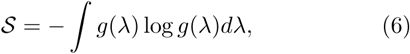

Where 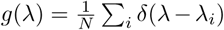 is the probabilistic density function of eigenvalues. This definition can also be understood as the Kullback–Leibler divergence from a uniform distribution to the given eigenvalue distribution (see SM, Sec. 4.2). Notably, the maximum-entropy distribution, in the absence of other constraints, corresponds to a uniform distribution. In such a situation, the minimum eigengap 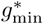 can be maximized. That is to say, to maximize the mutational robustness of the linear space 𝒱 spanned by eigenvectors, as shown in Fig. 2B, a uniform distribution is expected. Thus, for a system with many degrees of freedom, functional sensitivity and mutational robustness becomes two different optimization objectives that may be hard to be compatible.

### A. Entropy maximization

How can the dynamics of a protein molecule ensure that both functional sensitivity and mutational robustness are maintained? To resolve this problem, we introduced the entropy maximization approach to analyze the spacing problems of the eigenvalue distribution. Previously, the principle of maximum entropy [50] had already been applied in protein science. The applications include protein sequence evolution [51], structural constraint identification from correlated motions [52], residue interaction deduction [53], and protein folding [54]. In practice, the entropy maximization approach can handle multiple constraints, which introduce additional information into the system.

To determine the optimal eigenvalue distribution of protein systems, we consider fluctuation entropy as a constraint and maximize the spectrum entropy. From the physical perspective, it means that we fix the high functional sensitivity of a system, and then attempt to maximize the mutational robustness of the linear space spanned by all the eigenvectors. There are three constraints in this optimization process: (a) The spectrum distribution is normalized, i.e., ∫ *g*(*λ*)*dλ* = 1; (b) The total number of contacts in the proteins is constant, i.e., the trace of the Hessian matrix is fixed: ∫ *λg*(*λ*)*dλ* = *m*; (c) The fluctuation entropy is be fixed as a constraint 𝒞, i.e., ∫ log *λ* ·*g*(*λ*)*dλ* = 𝒞. Such a functional constraint ensures that during the evolution process, no matter how the interactions within proteins change with mutations, the fluctuation entropy will always be constant.

Take the three constraints as multipliers, we can obtain the Lagrangian expression:

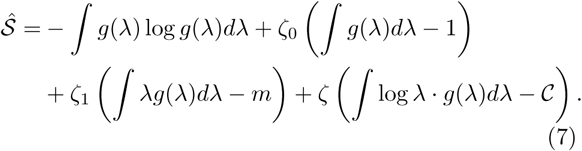

To find the distribution function *g*^*^(*λ*) that maximizes entropy 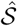 across all probability distributions, we require the following:

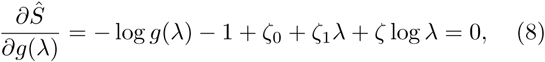

hence *g*^*^(*λ*) can be solved as:

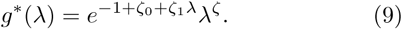

Such a spectrum shows a power-law distribution with an exponential cutoff. In the slow-mode limit, when *λ* ∼ 0, such a spectrum can be reduced into a power-law distribution: *g*(*λ*) ∼*λ*^*ζ*^, and the power-law exponent equals the Lagrangian multiplier *ζ*. In the fast-mode limit, the spectrum distribution will be dominated by the exponential function, which mainly results from the finite-size effect of the molecule. A similar eigenvalue distribution can be obtained by taking spectrum entropy as a constraint and maximizing the fluctuation entropy (listed in SM, Sec. 4.1).

Since proteins involve many degrees of freedom, such a distribution can achieve a great balance between sensitivity and robustness. For example, the external perturbations can easily trigger motions along the eigenvectors correspond to small eigenvalues. Meanwhile, other degrees of freedom can remain highly robust, reflecting that the energy landscape around the native basin of proteins is highly anisotropic, which is in line with previous molecular simulations [55–57]. Note that such anisotropy or heterogeneity is not encoded artificially into our model. Our definition of functional sensitivity and the mutational robustness of the whole subspace are both isotropic. Such anisotropy or heterogeneity emerges from the optimization process.

### B. The eigenvalue distribution of proteins

Remarkably, the spectra of natural proteins are in good accord with the above theoretical analysis. Here, we take three proteins (illustrated in Fig. 2C) with the same chain length (*N* = 200) but different structures as examples (more examples listed in SM, Sec. 5.1). Based on the vibration spectra (obtained by NMA) of proteins, the power-law coefficients can be obtained by fitting the eigenvalue distribution. Here, we consider the integrated eigenvalue distribution function 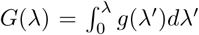. In the slow-mode limit, we have *g*(*λ*) ∼ *λ*^*ζ*^; thus, *G*(*λ*) ∼ *λ*^*ζ*+1^. As shown in Fig.2D, for the first 10 to 20 eigenvalues, *G*(*λ*) occurs in a line in the double logarithmic scale, clearly demonstrating power-law behavior. The slope of the log-log plot of *G*(*λ*) is used to determine the power-law coefficient *ζ* + 1. After fitting the first 20 distribution-based eigenvalues, for the three proteins in Fig. 2C, we have *ζ* = −0.17, 0.12, and 0.23.

Another approach for demonstrating power-law behavior is to obtain the rank-size distribution of the system. As discussed in the previous section, the magnitude of 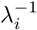 describes fluctuations along the direction of eigenvector 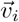. Because 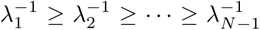, the rank-size relation of the eigenmodes is 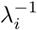 vs the rank (eigenvalue index) *i*. As shown in Fig. 2E, for slow-mode eigenvalues, the rank-size distribution shows a power law, known as Zipf’s law, i.e.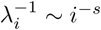, where *s* is the Zipf’s exponent. Fitting the first 20 eigenvalues in the spectra of the three proteins, we have *s* = 1.19, 0.88, and 0.78, which are all close to 1. Such a power-law decay quantitatively demonstrates that the low-frequency normal modes contribute significantly to the equilibrium fluctuations of proteins.

It is worth noting that the eigenvalue index *i* is proportional to the integrated eigenvalue distribution function,because 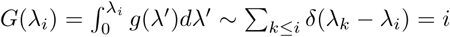. In the slow-mode limit, *G*(*λ*) ∼ *λ*^*ζ*+1^ ∼ *i* ∼ *λ*^1*/s*^. Thus, *ζ* = 1*/s* − 1 and measured *ζ* and *s* values are consistent with the relationship: For the three proteins in Fig. 2C, when *ζ* = − 0.17, 0.12, or 0.23, *s* = 1.19, 0.88, or 0.78. For most proteins, *s* ≈ 1, and *ζ* ≈ 0. Note that powerlaw coefficients *ζ* and *s* are closely related to the spectral dimension *d*_*s*_ [58] of the proteins: *d*_*s*_ = 2(*ζ* + 1) = 2*/s*, and *d*_*s*_ ≈ 2 *<* 3 (see SM, Sec. 4.3).

In Fig. 2F, the spectra of proteins with the same chain lengths (*N* = 200) are illustrated. Although the spectra of different proteins are varied, these spectra show similar behavior. At the logarithmic scale, the first 10 to 20 eigenvalues are roughly uniformly distributed, and act as a power-law distribution. The other eigenvalues are rather concentrated, demonstrating the characteristic scale of the exponential distribution. For comparison, the spectra of the Erdős-Rényi (ER) random network [59] and random geometric graphs (RGGs) with a similar number of nodes and edges [60] are also shown in Fig. 2F. The ER networks can be recognized as a simplification of extended or collapsed polymer systems with random correlations between the monomer units. The RGGs represent the random dense packing structures in the real space. In comparison with ER networks, the vibration spectrum of proteins has a notable power-law distribution in the slow mode. In contrast, the spectra of these proteins are very similar to those of RGGs. For most proteins, the scales of the eigenvalues (characterized by *λ*_*N*−1_*/λ*_1_) [61] lie between the spectra of 2-dimensional and 3-dimensional RGG. This result is robust for different kinds of weighted GNMs (see SM, Sec. 1.1) and is in line with those of previous studies, demonstrating that the structure of proteins can be described as the packing of amnio-acid residues in a fractal dimension between 2 and 3 [8, 58].

## IV. THE COMPATIBILITY OF FUNCTIONAL SENSITIVITY AND MUTATIONAL ROBUSTNESS: CONFIRMATION BY STRUCTURAL INFORMATION

Here, based on a large data set of protein structures (see Appendix C), we analyze the interplay between functional sensitivity and mutational robustness of proteins. For simplification, we mainly focus on the functional sensitivity and mutational robustness contributed by the first two normal modes (discussions on the other modes are listed in SM, Sec. 5.1). The fluctuation entropy contributed by the first two modes can be expressed as − log(*λ*_1_*λ*_2_); hence, we take 1/(*λ*_1_*λ*_2_) to be a descriptor of functional sensitivity. For mutational robustness, because *λ*_1_ and *λ*_2_ are the smallest eigenvalues in the system, to magnify the difference between them, we introduce 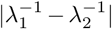 to describe the gap between the first and second eigenvalues.

Now, let us address the question regarding the physical meaning of the exponent *ζ*. We focus on a subset of proteins with similar chain lengths (180 ≤ *N <* 220). Although these proteins have similar sizes, their shapes and structures are different. For every protein, we fit the spectrum and obtain the value of *ζ*. Notably, the magnitude of *ζ* correlates with the functional sensitivity and mutational robustness of proteins. As shown in Fig. 3A, as *ζ* increases, the functional sensitivity described by 1/(*λ*_1_*λ*_2_) decreases. As shown in Fig. 3B, the mutational robustness described by 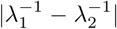 also decreases as *ζ* increases. Since both functional sensitivity and mutational robustness are negatively correlated with the coefficient *ζ*, as shown in Fig. 3C, for proteins with a given size, functional sensitivity is positively correlated with mutational robustness. This behavior resembles that of the three-bead system, as discussed in previous sections, which suggests the low-dimensionality of the dynamics of natural proteins.

**FIG. 3.**
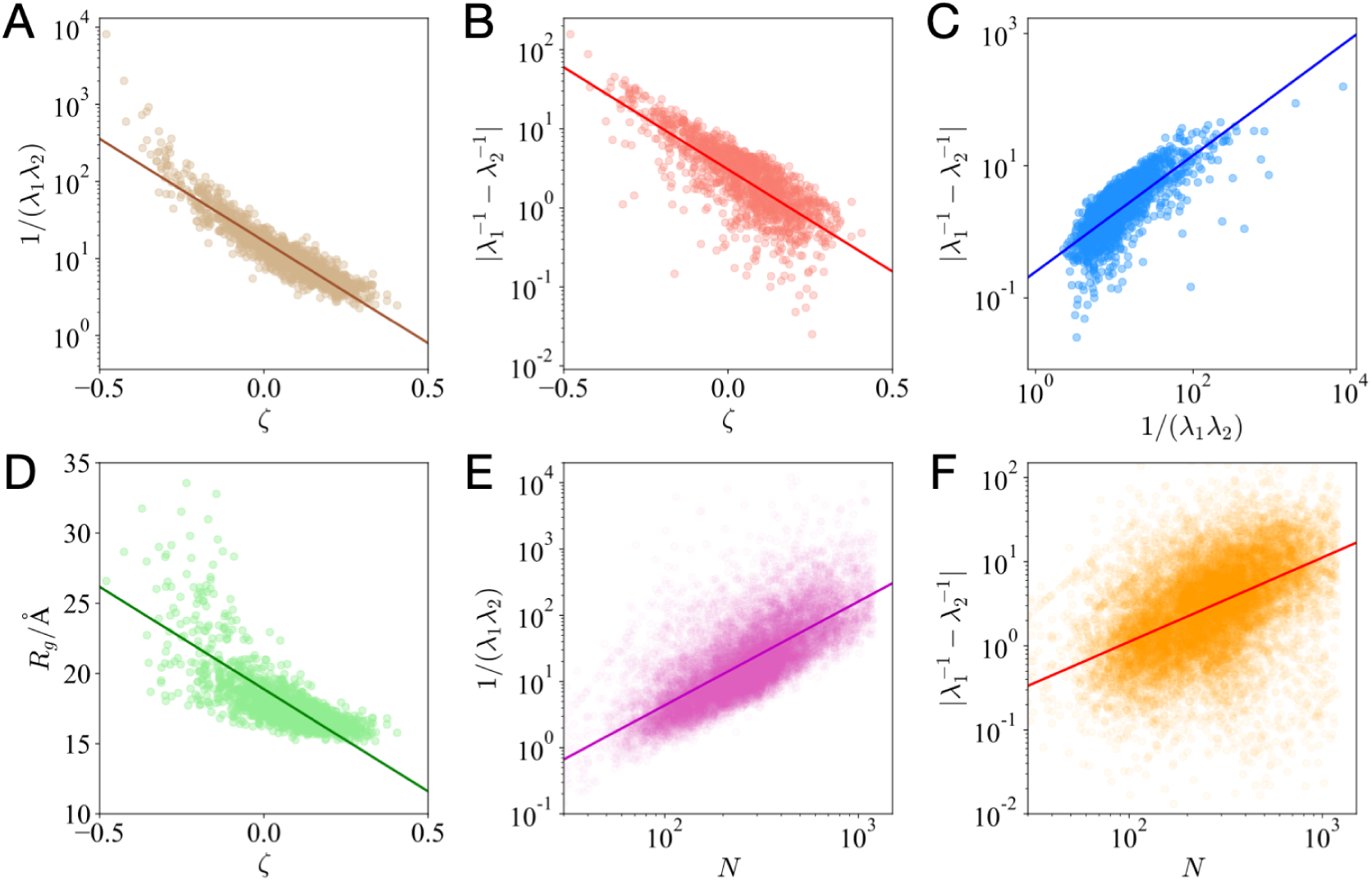
For proteins with similar sizes (chain length 180 *≤ N <* 220), the scattering plots (every data point denotes a protein) and trend lines of (A) functional sensitivity 1/(*λ*_1_*λ*_2_) vs the power-law coefficient *ζ*; (B) mutational robustness 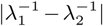 vs *ζ*; (C) mutational robustness 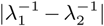 vs functional sensitivity 1/(*λ*_1_*λ*_2_); and (D) radius of gyration *R*_*g*_ vs *ζ*. For proteins with different sizes (30 *≤ N ≤* 1200), the scattering plots (every data point denotes for a protein) and trend lines of (E) functional sensitivity 1/(*λ*_1_*λ*_2_) vs chain length *N*, and (F) mutational robustness 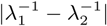 vs chain length *N* are shown.

The coefficient *ζ* is closely related to the protein structure. As shown in Fig. 3D, the radius of gyration *R*_*g*_ is negatively correlated with *ζ*. The three proteins illustrated in Fig. 2D can also support such an analysis. They have the same chain length, but their *R*_*g*_ and *ζ* are different. The chain length *N* per volume 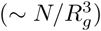 can act as an estimator of the packing density of proteins. A smaller *R*_*g*_ (larger *ζ*) indicates a higher packing density and lower flexibility of the molecule. On the contrary, a larger *R*_*g*_ (smaller *ζ*) implies a lower packing density and higher flexibility. The coefficient *ζ* is correlated with the fractal dimensions of proteins. When a protein is packed in lower spatial dimensions, there will be a smaller *ζ* and a larger *R*_*g*_. Besides, there will be a larger solvent-accessible surface area (SASA) [62] and higher modularity [63, 64], contributing to the large-scale motions and slow relaxations of the proteins (see SM, Sec. 5.2).

For natural proteins, both functional sensitivity and mutational robustness are scaled up as the protein size increases. As shown in Fig. 3E, as the chain length *N* increases, the magnitude of 1/(*λ*_1_*λ*_2_) also increases, showing that large proteins also have a higher functional sensitivity. The physical picture of this relationship is clear: small proteins tend to have a densely packed structure that ensures a stable folded state [65]; while for large proteins, slower vibration modes, which usually correspond with disordered loops or linkers, are demanded by the functionality. Meanwhile, as shown in Fig. 3F, as the protein size increases, the eigengap 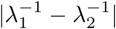 also increases, showing that slow vibration modes also exhibit a high mutational robustness. It is also understood that large proteins or protein complexes usually have multi-domain structures, which shows the high modularity of their residue contact networks [8]. The enhancement of the molecular flexibility contributes to inter-domain motions, which are usually highly robust. In all, the size dependence of functional sensitivity and mutational robustness suggests a general evolutionary constraint for proteins with different sizes.

## V. DISCUSSIONS

Plasticity and robustness are considered to be the two basic characteristics of biological systems [21–24]. They seem to be two opposite concepts at first glance; however, proteins can exhibit both the two characteristics. Native proteins are not only robust enough to tolerate random mutations, but also susceptible enough to sense environmental signals and perform large-scale conformational changes. It is suggested that a hidden link evolves between sensitivity/robustness to external perturbations and to mutations [23]. To reveal the relationship between functional sensitivity and mutational robustness, in this study, a general framework is presented to analyze the dynamics of a system under external and internal perturbations.

We demonstrate that for systems with low degrees of freedom, high functional sensitivity is equivalent to high mutational robustness. Biological systems, however, involve high degrees of freedom. Here, we show that these degrees of freedom are hierarchically organized, with power-law statistics of eigenvalues of relaxation dynamics. Larger conformational changes are governed by low-dimensional slow modes. Recent reports have shown that the dimensions of biologically relevant modes are reduced through evolution [66, 67]. Such evolutionary dimensional reduction was also observed in the phenotypic dynamics of proteins [28, 29, 68]. Such a property also concerns with “sloppiness” [69–71]. Systems are usually sensitive to perturbations along a few stiff dimensions that correspond with the functional dynamics and highly robust to mutations along many other sloppy dimensions. Both functional sensitivity and mutational robustness can be achieved with these sloppy systems. From the perspective of statistical inference, such systems can have both high sensitivity of the variables and high robustness in the estimation of the inferred parameters [72]. The low-dimensional dynamics have a slower timescale in general. These slow-mode motions usually overlap significantly with displacement during functional motions. These functional motions usually involve relative movements of large protein subunits or cooperative conformational changes in entire proteins. They confer a high functional sensitivity; a small perturbation can lead to motions along the directions of these modes. Previous research has shown that weak interactions at the interfaces of domains enable proteins to exhibit large-amplitude conformational changes [73–75], contributing to the functional sensitivity of the molecule. Meanwhile, these functional slow modes are also highly robust to mutations. Modularized structures in residue contact networks define gaps between the eigenvalues of the graph Laplacian [63, 64, 76], and contribute to mutational robustness (details in SM, Sec. 5.2). In practice, the gap between the first and second eigenvalues (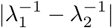 or log(*λ*_2_*/λ*_1_)) has been applied as an empirical descriptor of protein fitness [28, 29]. One theoretical interpretation is that when protein dynamics are low-dimensional, the maximization of the gap between the first two non-zero eigenvalues can contribute to the functional sensitivity and mutational robustness of a protein molecule.

For systems with high degrees of freedom, our results show that the compatibility between functional sensitivity and mutational robustness will lead to a power-law eigenvalue distribution in the spectrum, which is a sign of criticality. Our statistics for natural proteins reveal that small proteins prefer a densely packed structure to ensure a stable folded state, while large proteins have higher flexibility, which is required for their functionality. Such results reveal a universal principle in the evolution of proteins with different sizes and shed light on the design of functional proteins. Interestingly, staying at the critical point seems to be a common organizing principle of numerous varieties of biological systems [20, 77–82]: If a system is too disordered, it cannot stably exist or be reproduce; if it is too ordered, it cannot adapt or respond to environmental perturbations. Considering that proteins satisfy both functional sensitivity and mutational robustness, the theory we proposed here may explain why the proteins after evolution are near the critical points, and suggest one possible scenario for the evolution towards criticality.

## ACKNOWLEDGMENTS

We wish to thank Alexander S. Mikhailov, Holger Flechsig, Atsushi Kamimura, Takyuya U Sato, Lei-Han Tang, and Changsong Zhou for participating in stimulating discussions. This research was partially supported by a Grant-in-Aid for Scientific Research on Innovative Areas (17H06386), from the Ministry of Education, Culture, Sports, Science and Technology of Japan, and a Grant-in-Aid for Scientific Research (A)20H00123 from the Japanese Society for the Promotion of Science.

## Appendix A: GNM and Graph Laplacian

In Gaussian network model (GNM), the Hessian matrix ℋ then can be expressed as a Kirchhoff matrix that described residue contact topology [32, 33]. The entries in matrix ℋ are calculated as:

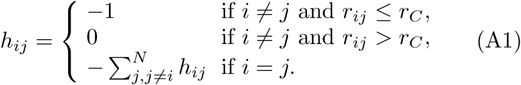

Here, *r*_*C*_ is the cut-off distance. In this study, we used *r*_*C*_ = 8.0Å. In practise, one can also introduce more accurate distance-dependent weights to describe the interaction strength between residue pairs. Additional discussions on the weighted GNM are listed in SM (Sec. 1.1).

According to Eq. A1, matrix ℋ can be expressed as ℋ = 𝒟 − A, in which 𝒟 is a diagonal matrix that describes the degrees of the nodes, and A is the adjacency matrix of the elastic network. For the matrix ℋ, the trace ∑_*i*_ ℋ_*ii*_ = ∑_*i*_ *λ*_*i*_ = 2*E*, in which *E* denotes the total number of edges in the elastic network. In a protein molecule, the number of edges, i.e. *E* indicates the total number of covalent and non-covalent residue contacts. Thus, *E* can work as an estimator of the energy of the native state.

## Appendix B: Correlated Motion

The correlated motion of native proteins can be described using the covariance matrix *C*, in which the matrix element 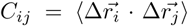 [34–36]. Experiments and molecular simulations can provide information on the correlated motions of the proteins. Based on the covariance matrix, by conducting principal component analysis (PCA), the eigenvalues and the eigenvectors describing the proteins’ dynamical modes can be obtained. It is observed that the low-frequency normal modes predicted by elastic network models can well match the eigenmodes given by PCA. In the GNM, the covariance matrix *C* is proportional to the pseudoinverse of the Hessian matrix 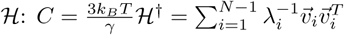 [17, 25, 32].

## Appendix C: Dataset

Our dataset contains 12954 proteins selected from the Protein Data Bank (PDB) [83]. The structures of these proteins were all determined via high-resolution X-ray diffraction (≤ 2.0Å). No DNA, RNA, or hybrid structures of proteins are included in the dataset, and the chain lengths are 30 ≤ *N* ≤ 1200. In our protein dataset, every two proteins share less than 30% sequence similarity. The PDB codes of all the proteins in our dataset are listed in the SM (Sec. 6.3).

