## Supplementary Materials for "Functional Sensitivity and Mutational Robustness of Proteins"

#### 1 Additional Discussions on Elastic Network Models

##### 1.1 The weighted Gaussian Network Model

In the Main Text, we had conducted our analysis based on the simplest version of the Gaussian network model (GNM). Our analysis can also be generalized to other versions of elastic network models.

**Original GNM.** Before we introduce the generalized model, let us first consider two nodes ($p$ and $q$, $p \neq q$) connected by a uniform spring. The distance between the two nodes is $r_{pq}$, while at the zero-potential-energy point, the distance is $r^0_{pq}$. The potential energy of such a bond would be: $V_{pq} = \frac{\gamma}{2}(r_{pq} - r^0_{pq})^2 = \frac{\gamma}{2}(\Delta\vec{r}_p - \Delta\vec{r}_q)^2$, in which $\gamma$ is the spring constant. In GNM, since the three-dimensional directional motions of the residues are simplified as homogeneous, we can dismiss the vector notions for $\Delta\vec{r}_p$ and $\Delta\vec{r}_q$. Then, the potential energy of such a bond can be written as:

$$V_{pq} = \frac{\gamma}{2}(\Delta r_p - \Delta r_q)^2 = \frac{\gamma}{2}\left(\Delta r_p^2 + \Delta r_q^2 \underbrace{-2\Delta r_p \Delta r_q}_{=-\Delta r_p\Delta r_q - \Delta r_q \Delta r_p}\right), \tag{S1}$$

which is a quadratic form. Note that the elements of the Hessian matrix $\mathcal{H}$ can be expressed as: $h_{ij} = \frac{\partial^2 V(\Delta r)}{\partial \Delta r_i \partial \Delta r_j}$. According to Eq.S1, in the elastic network, every bond $(p-q)$ will contribute to the off-diagonal elements $h_{pq}$ and $h_{qp}$ with $-1$, and the diagonal elements $h_{pp}$ and $h_{qq}$ with 1. By summing up the potential energy of all the bonds, the entries of matrix $\mathcal{H}$ can be calculated as:

$$h_{ij} = \begin{cases} -1 & \text{if } i \neq j \text{ and } r_{ij} \leq r_C, \\ 0 & \text{if } i \neq j \text{ and } r_{ij} > r_C, \\ -\sum_{j,j\neq i}^{N} h_{ij} & \text{if } i = j. \end{cases} \tag{S2}$$

**Weighted GNM.** When we consider different weights ($w_{ij}$) for different bonds, then the Hessian matrix $\mathcal{H}$ can be constructed as:

$$h_{ij} = \begin{cases} -w_{ij}, & \text{if } i \neq j, \quad \text{(S3)} \\ \sum_{j,i\neq j} w_{ij}, & \text{if } i = j. \quad \text{(S4)} \end{cases}$$

In applications, by introducing distance-dependent force constants, enhanced models were suggested to have better predictions on the B-factors or protein dynamics. Among the enhanced elastic network models, here in this Supplementary Material, we select the Harmonic C$_\alpha$ potential model (HCA) [1, 2] and the parameter-free Gaussian network model (pfGNM) [3] to compare with the simplest form of GNM.

**HCA Model.** In HCA model, the force constant $w_{ij}$ between residue $i$ and $j$ is defined as:

$$w_{ij} = \begin{cases} a_1 r_{ij}^0 - b, & \text{if } r_{ij}^0 < c \quad \text{(S5)} \\ a_2 (r_{ij}^0)^{-6}, & \text{if } r_{ij}^0 \geq c \quad \text{(S6)} \end{cases}$$

where $r_{ij}^0$ denotes the equlibrium distance between residue $i$ and $j$, and parameter $c$ is set to 4Å so that the interactions between two sequential-neighboring $C_\alpha$ atoms are considered to be different from other interactions. Other parameters ($a_1$, $a_2$ and $b$) are fitted from experimental data.

**Parameter-free GNM (pfGNM).** In pfGNM, the force constants follows the inverse-square of the equilibrium distance between the interacting nodes, that is:

$$w_{ij} = (r_{ij}^0)^{-2}. \quad \text{(S7)}$$

In the computation, one can also introduce other decaying exponent $p$, so that $w_{ij} = (r_{ij}^0)^{-p}$. When $p = 6$, then the model would be very close to HCA model.

**The Normalization of the Eigenvalues.** To compare the spectrum given by different weighted GNMs, the normalization of eigenvalues becomes an important issue. For example, for two different weighted GNM (model A and B), if the weight $w_{ij}^{(A)} = 2w_{ij}^{(B)}$, then the eigenvalues predicted by the two model should have $\lambda_i^{(A)} = 2\lambda_i^{(B)}$. In fact, these two models are effectively equivalent. To resolve this problem, there are usually two different ways to normalize the eigenvalues: (1) normalize by the mean value: $\hat{\lambda}_i = N\lambda_i / \sum_i \lambda_i$; or (2) normalize by the first (smallest) non-zero eigenvalue $\lambda_1$: $\tilde{\lambda}_i = \lambda_i/\lambda_1$.

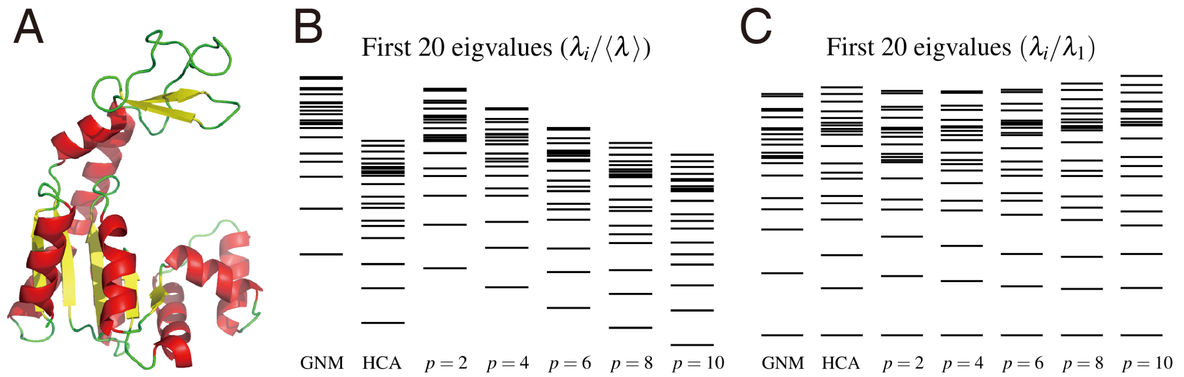


Figure S1: The spectrum of a protein with different types of weighted GNMs. (A) The cartoon illustration of the open-state adenylate kinase (PDB code: 4ake). (B) The first 20 eigenvalues of the protein with different weighted GNMs. The eigenvalues are normalized by their mean value ($\hat{\lambda}_i = N\lambda_i / \sum_i \lambda_i$) and shown in log-scale. (C) The first 20 eigenvalues of the protein with different weighted GNMs. The eigenvalues are normalized with the first non-zero eigenvalue ($\lambda_i/\lambda_1$) and shown in log-scale.

**Comparison of Different Models.** Here, we take the open-state adenylate kinase (ADK, Fig.S1A) as an example to show the effects of different kinds of weighted GNM. As shown in Fig.S1B, if we normalize the eigenvalues by the mean value, the first nonzero eigenvalues predicted by different models vary a lot. If we consider interactions in a shorter range (larger $p$ in pfGNM), as shown in Fig.S1B, the first eigenvalues predicted would be smaller and *vice versa.* Such a result is easy to understand. With only short-range interactions, the contact network has higher modularity, and the slow-mode motions (correspond to the relative motions between different modules) become easier to activate. Indeed, it is observed that the models with a shorter interaction range can better capture the long-range correlations and other important information in the solvated dynamics of protein molecules [2, 4]. However, if we normalize the eigenvalues by $\lambda_1$, as shown in Fig.S1B, the spectra given by different models show high similarity. In this work, to characterize the distribution of the eigenvalues of proteins (Fig. 2F of the Main Text), we normalized the eigenvalues with $\lambda_1$. Thus, the conclusions drawn from these data are highly robust.

##### 1.2 The Shannon Entropy of the Multivariate Gaussian Distribution

In the Main Text, we defined the fluctuation entropy $S_F$ as the Shannon entropy (or differential entropy) of the multivariate Gaussian distribution $\phi(\Delta\vec{r})$. Here we give the derivation of Eq. (3) of the Main Text.

$$
\begin{aligned}
S_F(\Delta\vec{r}) &= -\int \phi(\Delta\vec{r}) \log \phi(\Delta\vec{r}) d\Delta\vec{r} \\
&= -\int \phi(\Delta\vec{r}) \log \left( \frac{1}{(2\pi)^{3N/2} (\det \mathcal{H})^{-3/2}} \right) d\Delta\vec{r} - \int \phi(\Delta\vec{r}) \log \left( e^{-\frac{\gamma}{2k_BT} \Delta\vec{r}^T \cdot \mathcal{H} \cdot \Delta\vec{r}} \right) d\Delta\vec{r} \\
&= \left( \frac{3N}{2} \log(2\pi) - \frac{3}{2} \log(\det \mathcal{H}) \right) \underbrace{\left( \int \phi(\Delta\vec{r}) d\Delta\vec{r} \right)}_{=1} + \frac{\gamma}{2k_BT} \int \phi(\Delta\vec{r}) \log \left( e^{\Delta\vec{r}^T \cdot \mathcal{H} \cdot \Delta\vec{r}} \right) d\Delta\vec{r} \\
&= \frac{3N}{2} \log(2\pi) - \frac{3}{2} \log(\det \mathcal{H}) + \frac{\gamma}{2k_BT} \int \phi(\Delta\vec{r}) \log \left( e^{\mathrm{Tr}[\mathcal{H} \Delta\vec{r} \cdot \Delta\vec{r}^T]} \right) d\Delta\vec{r} \\
&= \frac{3N}{2} \log(2\pi) - \frac{3}{2} \log(\det \mathcal{H}) + \frac{\gamma}{2k_BT} \log \exp \mathrm{Tr} \left[ \mathcal{H} \underbrace{\int \phi(\Delta\vec{r}) \Delta\vec{r} \cdot \Delta\vec{r}^T d\Delta\vec{r}}_{\text{The definition of covariance}} \right] \\
&= \frac{3N}{2} \log(2\pi) - \frac{3}{2} \log(\det \mathcal{H}) + \frac{\gamma}{2k_BT} \cdot \underbrace{\frac{3k_BT}{\gamma} \log \exp \mathrm{Tr} \left[ \mathcal{H} \mathcal{H}^{-1} \right]}_{C = \frac{3k_BT}{\gamma} \mathcal{H}^{-1}} \\
&= \frac{3N}{2} \log(2\pi) - \frac{3}{2} \log(\det \mathcal{H}) + \frac{3N \log e}{2} = 3N s_0 - \frac{3}{2} \log(\det \mathcal{H}) \\
&= 3N s_0 - \frac{3}{2} \log \prod_{i=1}^{N-1} \lambda_i = 3N s_0 - \frac{3}{2} \sum_{i=1}^{N-1} \log \lambda_i,
\end{aligned}
$$

in which $s_0 = \frac{1}{2} \cdot \log(2\pi e)$, and the constant coefficient $3N$ corresponds to the 3 degrees of freedom of all the $N$ nodes.

#### 2 Additional Discussions on Functional Sensitivity

In numerical computations, the condition number is usually introduced to measure the sensitivity of a function, say, how much would the errors or perturbations in the input lead to the error in the output. In this work, based on the Gaussian network model (GNM), the protein molecules are simplified as linear systems. Then, we can define a condition number to measure the sensitivity of a protein molecule. Such a definition is consistent with the fluctuation entropy as defined in the Main Text.

Here, as shown in Fig. S2, we take external force $\vec{F}$ (such as protein hydration, ligand binding, protein-protein interactions, or other perturbations from the environment) as the input, and take the molecular deformation $\Delta\vec{r}$ from the native state (molecular vibrations or conformational changes) as the output. In the linear regime, the relationship between the output $\Delta\vec{r}$ and the input $\vec{F}$ would be: $\vec{\Delta r} = \mathcal{H}^{\dagger}\vec{F}$, where $\mathcal{H}^{\dagger}$ denotes the pseudoinverse of the Hessian matrix $\mathcal{H}$. In a fluctuating environment, when external perturbations (input arguments) $\vec{f}$ act

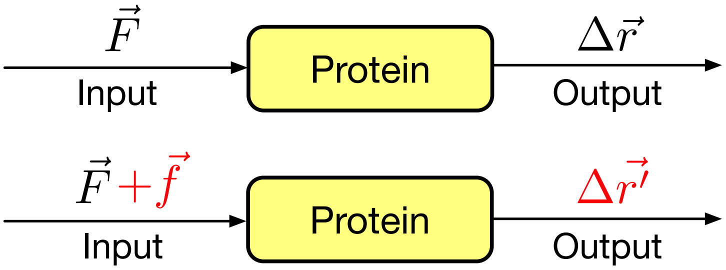


Figure S2: A protein molecule is modelled as a linear system. With the external force $\vec{F}$ as the input, the molecule would have deformation $\Delta\vec{r}$ as the output. When input arguments $\vec{f}$ is added to the molecule, then the output is modified as $\Delta\vec{r'}$.

on the molecule, the deformation (output errors) of the molecule changes from $\Delta\vec{r}$ to $\Delta\vec{r'}$. The relative change in the deformation (output) would be $||\Delta\vec{r'}-\Delta\vec{r}||/||\Delta\vec{r}||$ and the relative change in the external force (input) would be $||\vec{f}||/||\vec{F}||$. The ratio of the relative output ($\Delta\vec{r}$) error versus relative input ($\vec{F}$) error

$$c = \frac{\frac{||\Delta\vec{r'}-\Delta\vec{r}||}{||\Delta\vec{r}||}}{\frac{||\vec{f}||}{||\vec{F}||}} = \frac{||\mathcal{H}^{\dagger}\vec{f}||}{||\mathcal{H}^{\dagger}\vec{F}||}\cdot\frac{||\vec{F}||}{||\vec{f}||} \tag{S8}$$

We define sensitivity of a protein molecule as the maximum value $c^*$ of the ratio $c$, which is given by

$$c^* = \frac{\max||\mathcal{H}^{\dagger}\vec{f}||\cdot||\vec{F}||}{\min||\mathcal{H}^{\dagger}\vec{F}||\cdot||\vec{F}||} = \frac{\lambda_{\max}}{\lambda_{\min}} = \frac{\lambda_{N-1}}{\lambda_1}. \tag{S9}$$

The value of $c^*$ describes how much would the perturbations in external forces lead to the changes in the deformation of a protein molecule. Note that the value $c^*$ can be understood as a generalization of condition number, which is widely applied in the measurement of the intrinsic sensitivity of linearized dynamics. An ill-conditioned protein molecule can exhibit large conformational changes with low energy costs, which is highly beneficial in performing its functional dynamics. An ill-conditioned network structure corresponds to a protein molecule shows high sensitivity under external perturbations.

Note that the condition number $c^*$ has its limitations: the optimization towards the maximum condition number can only affect the largest and the smallest eigenvalue. Therefore, in the Main Text, we introduce fluctuation entropy to quantify the functional sensitivity of the molecule. According to Eq.(3) in the Main Text, the fluctuation entropy $S_F$ is a convex function of eigenvalue $\lambda_j$'s. When we fix the total number of contact (total number of edges) as $E$, the fluctuation entropy $S_F$ and the condition number $c^*$ will be two correlated descriptors of sensitivity: according to Jensen's inequality, the fluctuation entropy reaches the minimum when all the eigenvalues take the same value. In such a situation, the condition number $c^*$ also reaches minimum ($c^* = 1$). The fluctuation entropy reaches maximum when the first $N-2$ vibration modes have the corresponding eigenvalues that ($\lambda_1 \approx \lambda_2 \approx \cdots \approx \lambda_{N-2} \approx 0$) and the fastest mode correspond to $\lambda_{N-1} \approx 2E$. In such a situation, the condition number $c^* \to \infty$.

### 3 Additional Discussions on Mutational Robustness

#### 3.1 An Illustrative Model

In the Main Text, our analysis of the mutational robustness is mainly based on the elastic networks' spectral properties. Here, we take a specific kind of mutation (residue substitution) as an example to show how our method can be applied in the analysis of the mutational robustness of a protein. For simplification, it is assumed in our model that the mutations would only have minor effects on the structure of the native state, that is, the mutations at a single site will not change the structure of the elastic network significantly.

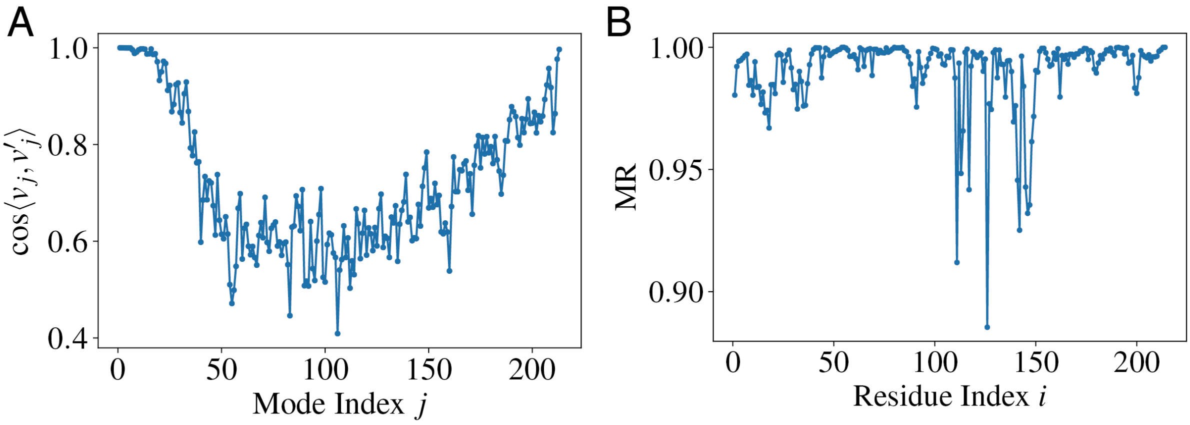


Figure S3: The mutational robustness of ADK molecule. (A) The mutational robustness (defined as the cosine of the intersection angle between the perturb $v'_j$ and unperturbed $v_j$) versus mode index $j$. (B) The mutational robustness $\mathrm{MR}_i$ for residue substitution at different sites.

Here, we take the ADK molecule (Fig. S1A) as an illustrative example. There are 214 amino acid in the chain of the ADK molecule. Let us consider a residue substitution at site $p$ (as illustrated in Fig. 1B of the Main Text). After the mutation, the interactions between residue $p$ and other neighboring residues will be weakened. In such a case, the perturbation matrix $M$ would have the entries $M_{pj} = M_{jp} = -h_{pj}$ for all $j \neq p$, $M_{pp} = -h_{pp}$, and $M_{jj} = h_{pj}$, where $h_{pq}$'s are the entries of the original Hessian $\mathcal{H}$. Then, we take $\epsilon = 0.5$, for the perturbed Hessian matrix $\mathcal{H}' = \mathcal{H} + \epsilon M$, the $j$-th eigenvector will changed from $\vec{v}_j$ to $\vec{v}'_j$. For simplification, in the following discussions, we omit the notations of vectors and write $\vec{v}_j$ as $v_j$.

By randomly introducing amino acid substitution at the sequence, we measured the average cosine of the intersection angles $\cos\langle v_j, v'_j\rangle$ for all the nonzero modes. As shown in Fig.S3A, after the mutation, there are about 20 slow modes, and 2 fast modes can approximately keep its direction of the motion ($\cos\langle v_j, v'_j\rangle > 0.95$), showing high mutational robustness. In fact, for these eigenvectors, the corresponding eigenvalues (for matrix $\mathcal{H}$ or its pseudoinverse) have large gaps with neighboring eigenvalues. Such a result is also in line with previous studies [5, 6]. These robust slow modes are correlated with the functional motions, and the fast modes correspond to the local constraints in a molecule.

Even the first 20 slow modes show relatively high mutational robustness under residue substitution, the mutations at different sites may lead to different functional consequences. To quantify such an effect, we can introduce the mutational robustness for every amino acid residue in a protein sequence. For example, after the mutation at residue $i$, the first $m$ modes change consequently ($v_j \to v_j'^{(i)}$), and for each mode, we can calculate the cosine of the intersection angle $\cos\langle v_j, v_j'^{(i)}\rangle$. Then, we define the average cosine of the first $m$ modes as mutational robustness of residue $i$:

$$\mathrm{MR}_i = \frac{1}{m}\sum_{j=1}^{m} |\cos\langle v_j, v_j'^{(i)}\rangle|. \tag{S10}$$

A large $\mathrm{MR}_i$ value correspond to high mutational robustness at site $i$. As shown in Fig.S3B, the residues 111, 113, 117, 126, 141, 142, 145, 146, and 147 are predicted to have lowest robustness during the mutation ($\mathrm{MR}_i < 0.95$), whereas others keep high mutational robustness. In fact, these residues are among the hot residues during the conformational changes (from open state to closed state) [7].

##### 3.2 Mutational robustness based on the intersection angle

In the Main Text, to quantify the mutational robustness of the $j$-th eigenvector $v_j$, we introduced the square deviation $|\Delta v_j|^2$ to measure the "distance" from the original $v_j$ to $v'_j$. Another way to quantify the robustness of an eigenvector is to estimate the intersection angle $\langle v_j, v'_j\rangle$ between the unperturbed and perturbed eigenvectors. It is worth noting that, with other definitions of the "distance" between two eigenvectors, our analysis based on the eigengaps can also hold.

When Hessian matrix changes from $\mathcal{H}$ to $\mathcal{H}'$, where $\mathcal{H}' = \mathcal{H} + \epsilon M$. According to Davis-Kahan Theorem [8, 9], the difference between the unperturbed eigenvector $v_j$ and the perturbed $v'_j$

$$\sin\langle v_j, v'_j\rangle \leq \frac{2 \cdot \epsilon ||M||_{\mathrm{op}}}{\min(\lambda_{j+1} - \lambda_j, \lambda_j - \lambda_{j-1})}, \tag{S11}$$

in which $||M||_{\mathrm{op}}$ is the operator norm of matrix $M$, which is defined as the largest singular value of matrix.

As shown in Eq.S11, the robustness of the $j$-th eigenvector is determined by the magnitude of the perturbation $\epsilon$, the operator norm $||M||_{\mathrm{op}}$ and the minimum gap between neighboring eigenvalues. Here, the magnitude of perturbation $\epsilon$ can be recognized as a constant, and for adding or removing a contact, there will be $||M||_{\mathrm{op}} = 1$. In such a situation, the robustness of the eigenvector $v_j$ is only determined by the gaps between the neighboring eigenvalues $\lambda_j - \lambda_{j-1}$ and $\lambda_{j+1} - \lambda_j$. Such a result is also in line with the classical perturbation theory as discussed in the Main Text.

##### 3.3 The inverse of the eigenvalues

For the low-frequency modes with $\lambda_i \approx 0$, the eigengaps $|\lambda_i - \lambda_{i-1}|$ are too small, according to Davis-Kahan theorem, there will always be a very high upper bound for the intersection angle. To have an accurate estimation of the mutational robustness, one shall use the eigengaps in the psedudoinverse of the matrix $\mathcal{H}$ (which is proportional to covariance matrix) to estimate how much would such a mutation affect the covariance matrix $C$ ($\sim \mathcal{H}^\dagger$). We have

$$\begin{aligned}(\mathcal{H}')^\dagger &= \Big(\mathcal{H}(I + \epsilon\mathcal{H}^\dagger M)\Big)^\dagger = (I + \epsilon\mathcal{H}^\dagger N)^\dagger \cdot \mathcal{H}^\dagger \\ &\approx \mathcal{H}^\dagger - \epsilon\mathcal{H}^\dagger M \mathcal{H}^\dagger.\end{aligned} \tag{S12}$$

Then, according to Davis-Kahan theorem, we have

$$\sin\langle v_j, v'_j\rangle \leq \frac{2 \cdot \epsilon ||\mathcal{H}^\dagger M \mathcal{H}^\dagger||_{\mathrm{op}}}{\min(\lambda_{j+1}^{-1} - \lambda_j^{-1}, \lambda_j^{-1} - \lambda_{j-1}^{-1})}. \tag{S13}$$

Eq.S13 shows that the mutational robustness of $v_j$ is bounded by the eigengaps in the $\mathcal{H}^\dagger$. Therefore in the Main Text, we introduce $|\lambda_1^{-1} - \lambda_2^{-1}|$ to describe the robustness of the $v_1$.

##### 3.4 Distance between linear subspaces

The Davis–Kahan theorem can also be used in the perturbation analysis to bound the distance between linear subspaces. For $N \times N$ symmetric real matrix $\mathcal{H}$ and $\mathcal{H}'$, their eigenvectors are $v_1, v_2, \cdots v_N$ and $v'_1, v'_2, \cdots v'_N$, respectively. As discussed in Ref. [9], let us consider the distance between the two subspace $V = (v_r, v_{r+1}, \cdots, v_s) \in \mathbb{R}^{N\times d}$ and $\hat{V} = (v'_r, v'_{r+1}, \cdots, v'_s) \in \mathbb{R}^{N\times d}$ $(1 \leq r \leq s \leq N)$.

To calculate the "distance" between the two subspace, we compare the difference in the principal angles between the two subspace. To determine the angle, we calculate the inner product of the two subspaces $\hat{V}^T V$, and there are $d$ singular values $(\sigma_1, \sigma_2, \cdots \sigma_d)$ for matrix $\hat{V}^T V$. Since both $V$ and $\hat{V}$ have orthonormal columns, these singular values reflect the $d$

principal angles between their column spaces, which are given by $\cos^{-1}\sigma_1, \cos^{-1}\sigma_2, \cdots \cos^{-1}\sigma_d$. Then, we define the $\Theta(V, \hat{V})$ as a $d \times d$ diagonal matrix in which $\Theta_{jj} = \cos^{-1}\sigma_j$, and $1 \leq j \leq N$.

According to the useful variant of the Davis-Kahan theorem [9], the distance $\Theta(V, \hat{V})$ between the two $d-$dimensional subspace $V$ and $\hat{V}$ would be bounded as:

$$||\sin\Theta(\hat{V}, V)||_{\mathrm{F}} \leq \frac{2 \cdot \min(d^{1/2}||\hat{A} - A||_{\mathrm{op}}, ||\hat{A} - A||_{\mathrm{F}})}{\min(\lambda_{r-1} - \lambda_r, \lambda_s - \lambda_{s+1})}, \tag{S14}$$

in which $||\cdot||_F$ is the Frobenius Norm of the matrix, which is defined as $||A||_{\mathrm{F}} = \sqrt{\mathrm{Tr}(A^T A)} = \sqrt{\sum_{i=1}^{M}\sum_{j=1}^{N}|a_{ij}|^2} = \sqrt{\sum_{i=1}^{r}\sigma_i^2}$. Eq.S14 shows that the robustness of the eigenvectors against matrix element perturbation is determined by the eigengaps of the matrix. For every eigenvalue $\lambda_j$ of matrix $A$, there are two eigengaps: $\lambda_{j-1} - \lambda_j$ and $\lambda_j - \lambda_{j+1}$ (For $j = 1$, here we define $\lambda_0 = -\infty$, and for $j = N$, we define $\lambda_{N+1} = \infty$.) These eigengaps work as the intrinsic attributes which determine the robustness of a matrix under perturbation. As illustrated in Fig.S4, the distance between the perturbed and unperturbed subspace is bounded by the eigengaps at the boundary of the (unperturbed) subspace.

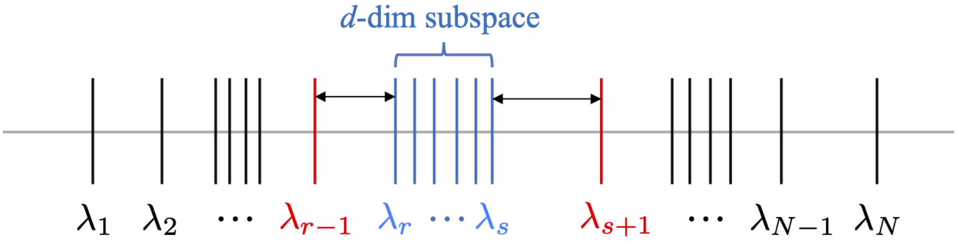


Figure S4: The robustness of the subspace is determined by the eigengaps. The distance between the perturbed and unperturbed subspace can be determined by the eigengaps at the boundary of the subspace.

We can also define the distance between two linear spaces based on the Chebyshev distance. In mathematics, the Chebyshev distance between two vectors is defined as the maximum of their differences along any coordinate dimension. Therefore, such a metric is also called the "maximum metric". One can also generalize such a metric to quantify the distance between two matrices (linear space) $V$ and $\hat{V}$, that is, the distance between two matrices is defined as the maximum of their differences along any column (the eigenvector with the same ordinal value). Such a metric is intuitive, to measure the distance between two linear spaces, one should first determine which column has changed the most, and then measure how much it changes for such a column. With such a metric, the distance between two eigenspaces is defined as the maximum distance between the eigenvectors (or columns) with the same ordinal value from the two eigenspaces.

Then, one can define the maximum Euclidean distance between the two eigenspaces as

$$\max_{1 \leq i \leq d} ||v'_i - v_i||_2,$$

that is, the distance between two eigenspaces is defined as the maximum Euclidean distance between the two eigenvectors with the same ordinal value from the two eigenspaces.

Or one can define the maximum Chebyshev distance between the two eigenspaces as

$$\max_{1 \leq i \leq d}\left(\max_{1 \leq j \leq N}|v'_{ij} - v_{ij}|\right),$$

that is, the distance between two eigenspaces is defined as the maximum Chebyshev distance between the two eigenvectors with the same ordinal value from the two eigenspaces.

In brief, for the eigenspace which has larger eigengaps between neighboring eigenvalues, such an eigenspace would be more robust under perturbation. Therefore, to maximize the robustness of the eigenspace, one should maximize the minimum eigengap between neighboring eigenvalues.

##### 3.5 Uniform eigenvalue distribution relates to the highest eigenspace robustness

In this subsection, we prove that, when the magnitude of the perturbation (the norm of the perturbation matrix) is fixed, a uniform eigenvalue distribution gives the highest eigenspace robustness. Consider the eigenspace $V = (v_1, v_2, v_3, \cdots, v_d) \in \mathbb{R}^{N\times d}$ correspond with non-negative eigenvalues $\{\lambda_1, \lambda_2, \lambda_3, \cdots \lambda_d\}$, and $0 \le \lambda_1 \le \lambda_2 \le \lambda_3 \le \cdots \le \lambda_d$. With a fixed summation of eigenvalues (i.e., fix $\sum_i \lambda_i = S > 0$), to maximize the minimum eigengaps $g_{\min}$, the spectrum $\{\lambda_1, \lambda_2, \lambda_3, \cdots \lambda_d\}$ should be an arithmetic series, $\lambda_1 = 0$ with a constant difference, i.e., $\lambda_i = c(i-1)$ with $c = \frac{2s}{d(d-1)}$.

Assume that there is a non-negative sequence $\{\lambda'_1, \lambda'_2, \lambda'_3, \cdots \lambda'_d\}$ other than the arithmetic series can maximize the minimum eigengaps with fix summation $S$, and the minimum gap $g^*$ could be found between the $j$-th and $(j-1)$-th eigenvalue, then we have $\lambda_j - \lambda_{j-1} = g^* > g_{\min}$, and $\lambda_j - \lambda_{j-1} < \lambda_i - \lambda_{i-1}$ for $j \neq i$. In such a situation, the summation of all the eigenvalue

$$S = \sum_{i=1}^{d} \lambda'_i \ge d\lambda'_1 + \frac{d(d-1)g^*}{2} > \frac{d(d-1)g^*}{2} > \frac{d(d-1)g_{\min}}{2} = S,$$

which lead to $S > S$. Therefore, the assumption is not true. Thus, the spectrum which maximizes the minimum eigengaps would be an arithmetic series. In the Main Text, we introduce the Shannon entropy of the spectrum to evaluate the mutational robustness of the system, because the maximum-entropy distribution (without other constraints) would be the uniform distribution that can maximize the minimum gaps between neighboring eigenvalues.

#### 4 Remarks on the Entropy Maximization

##### 4.1 Maximizing fluctuation Entropy

In the Main Text, by taking fluctuation entropy as a constraint, we maximize the spectrum entropy of the system. Similar distribution can also be obtained by taking spectrum entropy as a constraint and maximize the fluctuation entropy. In such an optimization process, there are three constraints:

(a) The spectrum distribution should be normalized, i.e., $\int g(\lambda)d\lambda = 1$;

(b) The total number of contacts in the proteins is a constant, i.e., to fix the trace of the Hessian matrix $\int \lambda g(\lambda)d\lambda = m$;

(c) Robustness constraint: $\int g(\lambda)\log g(\lambda)d\lambda = \mathcal{C}'$.

Take the three constraints as Lagrangian multiplier, then we can obtain the Lagrangian expression:

$$\hat{\mathcal{S}} = -\int \log\lambda \cdot g(\lambda)d\lambda + \eta_0\left(\int g(\lambda)d\lambda - 1\right) + \eta_1\left(\int \lambda g(\lambda)d\lambda - m\right) + \eta\left(\int g(\lambda)\log g(\lambda)d\lambda - \mathcal{C}'\right). \quad \text{(S15)}$$

To find the distribution function $g^*(\lambda)$ that maximize entropy $\hat{\mathcal{S}}$ across all probability distributions, we require that:

$$\frac{\partial\hat{\mathcal{S}}}{\partial g(\lambda)} = -\log\lambda + \eta_0 + \eta_1\lambda + \eta(\log g(\lambda) + 1) = 0, \quad \text{(S16)}$$

hence $g^*(\lambda)$ can be solved as:

$$g^*(\lambda) = e^{-1-\hat{\eta}_0-\hat{\eta}_1\lambda}\lambda^{1/\eta}, \tag{S17}$$

in which $\hat{\eta}_0 = \eta_0/\eta$ and $\hat{\eta}_1 = \eta_1/\eta$. Such a spectrum distribution is also a power-law distribution with an exponential cutoff, which agrees with the form obtained in the Main Text. In the slow-mode limit, the power-law exponent equals $1/\eta$.

#### 4.2 Kullback-Leibler divergence

In the Main Text, we mentioned that the spectrum entropy $S$ can be understood as the Kullback-Leibler divergence from the uniform distribution. Here is the proof.

Suppose $q(x)$ is a uniform distribution ($q(x) = c$ for $x$ defined on the interval $X$). For a given distribution $p(x)$, to minimize the K-L divergence from $q(x)$ to $p(x)$, we have

$$\begin{aligned} D_{KL}(p||q) &= \int_{x\in X} p(x)\log p(x)dx - \int_{x\in X} p(x)\log q(x)dx \\ &= -S - \int_{x\in X} p(x)\log c dx = -S - \log c\cdot ||X||, \end{aligned} \tag{S18}$$

in which $||X||$ is the measure of interval $X$, and it is a constant. Thus, minimizing the K-L divergence from given distribution $p(x)$ to uniform distribution $q(x)$ is equivalent with the maximization of spectrum entropy $S$.

#### 4.3 The eigenvalue distribution and the spectral dimension

In the Main Text, we take three proteins with the same chain length ($N = 200$) as examples to fit the eigenvalue distribution and obtain different values of the power-law coefficient $\zeta$. Here, more proteins are taken for analysis. For proteins with the chain length $N = 200$, the scattering plots of power-law coefficient $\zeta$ vs radius of gyration $R_g$ are shown in Fig.S5A. The linear fitting (red line) shows the basic trend of these data: as $R_g$ increases, $\zeta$ decreases. To demonstrate the relation between protein structure and the coefficient $\zeta$, among the proteins listed in Fig.S5A, 9 proteins are selected, and their structures are illustrated in Fig.S5B. These proteins have the same chain length ($N = 200$), and they are ordered by their $R_g$. As shown in Fig.S5B, the protein with a positive $\zeta$ and a small $R_g$ usually have more secondary structures and are packed into a globular shape; while those proteins with a negative $\zeta$ and a large $R_g$ have more disordered segments or their shape deviate from a globule.

The value of $\zeta$ is closely related to the spectral dimension $d_s$ of the proteins. The spectral dimension $d_s$ is defined based on the eigenfrequency distribution [10]. In the slow-mode limit, when the eigenfrequency distribution obeys the power law: $f(\omega) \sim \omega^{d_s-1}$, in which $d_s$ denotes the spectral dimension of the protein, the integration of the eigenfrequency distribution $F(\omega) = \int_0^\omega f(\omega')d\omega' = \omega^{d_s}$. Note that in the discrete version, both $F(\omega_i)$ and $G(\lambda_i)$ correspond to the eigenvalue (or eigenfrequency) index $i$ of the system, so $i \sim \omega_i^{d_s} = \lambda_i^{\zeta+1}$. In normal mode analysis, the eigenvalue $\lambda_i$ is proportional to the square of the eigenfrequency $\omega_i^2$ [11]. Thus, we have $d_s = 2(\zeta+1)$. In our work, we take the cutoff distance $r_C = 8.0$Å. In such a situation, the spectral dimension $d_s \approx 2 < 3$, and $\zeta \approx 0$. As shown in Fig. S5C and D, both the spectral dimension $d_s$ and the power-law coefficient $\zeta$ are robust for the selection of different parameters (cutoff distance $r_C$). With a shorter cutoff distance, for example, $r_C = 6.5$Å (as selected in Ref.[10]), there will be a slightly smaller $d_s$ and $\zeta$ (because shorter cutoff distances corresponds to more slow modes [4]), but we still have $d_s \approx 2$ and $\zeta \approx 0$.

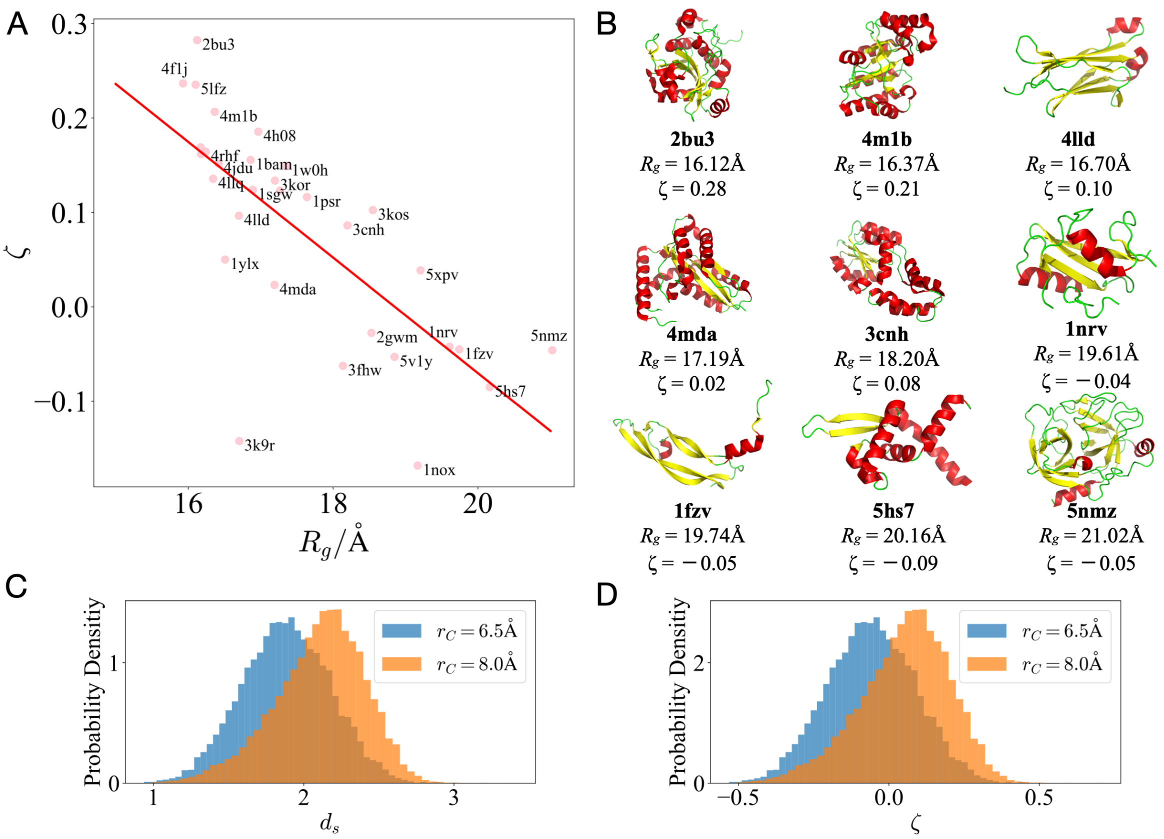


Figure S5: (A) For proteins with the same chain length ($N = 200$), the scattering plots and the linear fit of $\zeta$ vs $R_g$. Every data point denotes a protein. (B) The cartoon structure of 9 proteins with chain length $N = 200$. The PDB code, $R_g$, and $\zeta$ are also listed in the figure. (C) The histogram of the spectral dimension $d_s$ and (D) the histogram of the power-law coefficient $\zeta$ for all the proteins in our data set.

#### 5 Additional Discussions based on Protein Data

##### 5.1 The size-dependence of functional sensitivity and mutational robustness

In the Main Text, to uncover how the scaling coefficient $\zeta$ relates to the functional sensitivity and mutational robustness of a protein, we focus on a subset of proteins with similar chain lengths ($180 \leq N < 220$). The relation between functional sensitivity (or mutational robustness) and $\zeta$ also holds for proteins with different sizes. As shown in Fig. S6A, for protein at different sizes, as $\zeta$ increases, the functional sensitivity described by $1/(\lambda_1\lambda_2)$ decreases. As shown in Fig. S6B, for protein at different sizes, the mutational robustness described by $|\lambda_1^{-1} - \lambda_2^{-1}|$ also decreases as $\zeta$ increases. As shown in Fig. S6C, for protein at different sizes, functional sensitivity is still positively correlated with mutational robustness.

For natural proteins, the mutational robustness increase as the size of the protein increases. In the Main Text, we demonstrate that, as the size of the protein increases, the eigengap $|\lambda_1^{-1} - \lambda_2^{-1}|$ also increases, showing that the slow vibration modes also have high mutational robustness. Here, in Fig. S6D and Fig. S6E, the eigengap $|\lambda_2^{-1} - \lambda_3^{-1}|$ and $|\lambda_3^{-1} - \lambda_4^{-1}|$ also increase as the protein size increases. Such results suggest that, for real proteins, optimizing the mutational robustness of a specific slow mode may also contribute to the mutational robustness of other slow modes.

##### 5.2 The physical meaning of the power-law coefficient

To understand how the power-law coefficient $\zeta$ is related to the structure of the proteins, in the Main Text, we introduce the chain length per volume ($N/R_g^3$) to estimate the packing density of the amino-acid residues and discuss the relationship between the radius of gyration $R_g$ and $\zeta$. With only $R_g$ information, the protein is approximated as a perfect globule, in which all residue interactions are neglected. Such a description oversimplified the structure of the protein. Coefficient $\zeta$ can correlate with more detailed information on the structure and dynamics of the proteins.

###### 5.2.1 Solvent-accessible surface area (SASA)

In protein folding, the burial of hydrophobic amino-acid residues is a driving force. The flexibility of the proteins is closely related to the surface area buried within its fold [12]. If proteins are always folded into a perfect globule, the for proteins at similar sizes (similar chain length $N$), SASA $\sim R_g^2 \sim N^{2/3}$ should be a constant. For real proteins, however, their shape will deviate from the perfect globule, and their SASAs also vary. Here, we focus on a subset of proteins with similar chain lengths ($180 \leq N < 220$). As shown in Fig. S6F, as $\zeta$ decreases, the mean SASA increases, showing the deviation from the perfect globule. Such deviation from the globule will increase the SASA of the molecule, and confer high flexibility to the molecule. For real proteins, it is the solvation/hydration process [13] that plays an important role in the optimization of fluctuation entropy. Experiments and simulations demonstrated that the hydration structure's changes could contribute to protein flexibility and large-amplitude motions of the proteins [14, 15].

###### 5.2.2 Modularity

Previous research proved that the modularized structures in the residue contact network could contribute to the large-scale motions and slow relaxations of the proteins [4]. Coefficient $\zeta$ also correlates with the modularity of the residue contact networks. As shown in Fig. S6G, high modularity $Q$ correlate with small $\zeta$ (with negative values). As shown in Fig. S6H and Fig. S6I, for proteins at similar sizes, as modularity $Q$ of the system increases, both product of the

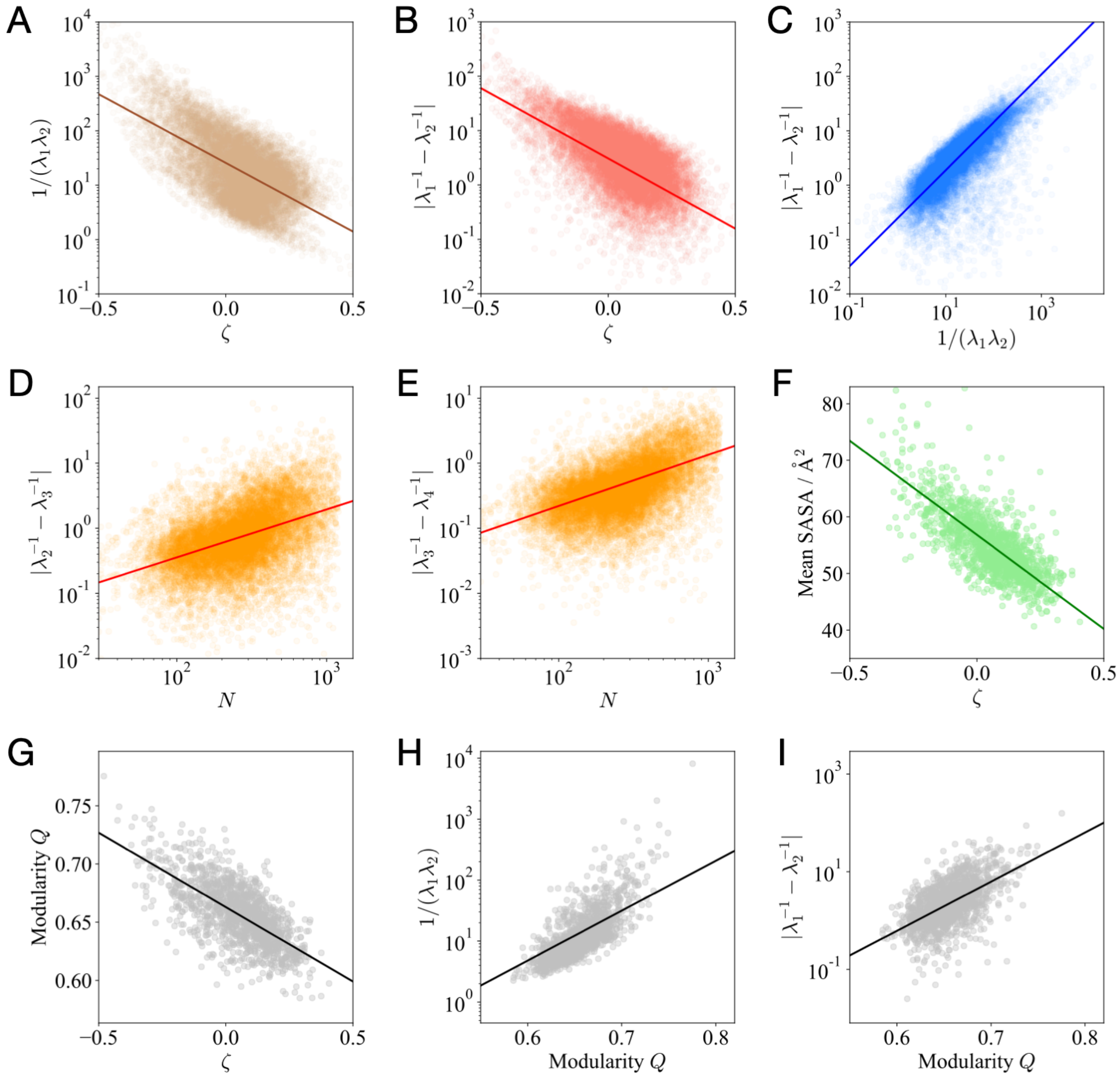


Figure S6: For protein of different sizes, the scattering plots (every data point denotes for a protein) and the trend lines of (A) functional sensitivity $1/(\lambda_1\lambda_2)$ vs the power-law coefficient $\zeta$; (B) mutational robustness $|\lambda_1^{-1} - \lambda_2^{-1}|$ vs $\zeta$; (C) mutational robustness $|\lambda_1^{-1} - \lambda_2^{-1}|$ vs functional sensitivity $1/(\lambda_1\lambda_2)$; (D) eigengap $|\lambda_2^{-1} - \lambda_3^{-1}|$ vs chain length $N$, and (E) $|\lambda_3^{-1} - \lambda_4^{-1}|$ vs chain length $N$. For proteins at a similar size (chain length $180 \leq N < 220$), the scattering plot of (F) mean solvent-accessible surface area (SASA) vs $\zeta$, (G) modularity $Q$ vs $\zeta$, (H) functional sensitivity $1/\lambda_1\lambda_2$ vs modularity $Q$ and (I) mutational robustness $\lambda_1^{-1} - \lambda_2^{-1}$ vs modularity $Q$.

eigenvalues $1/(\lambda_1\lambda_2)$ and the eigengap $|\lambda_1^{-1} - \lambda_2^{-1}|$ increase, demonstrating that modularized structures can contribute to both functional sensitivity and mutational robustness.

### 6 Dataset and Methods: Additional Discussions

#### 6.1 Solvent-accessible Surface Area (SASA)

The solvent-accessible surface area (SASA) defined as the surface area of a protein that is accessible to a water molecule. In this work, we introduced Shrake-Rupley algorithm [16] (embedded in Python Package "MDTraj" [17]) to measure the SASA of the protein molecule.

#### 6.2 Fiedler Vector and Modularity

The Fiedler vector of a graph, namely the eigenvector $v_1$ corresponding to the smallest nonzero eigenvalue $\lambda_1$ of the graph Laplacian matrix, plays an important role in spectral graph partitioning [18, 19]. Based on the sign of the corresponding vector entry, the Fiedler vector $v_1$ bisects the network into two communities. In the community detection of real-world networks, modularity $Q$ is defined as the fraction of the edges that fall within the given module minus the expected fraction when edges were distributed at random [20, 21]. Such a quantity is designed to quantify if a network can be easily divided into modules. For a network with $N$ nodes and $E$ edges, when the topology is described by the adjacency matrix $\mathcal{A}$, one can introduce the modularity matrix $\mathcal{B}$ with elements $\mathcal{B}_{ij} = \mathcal{A}_{ij} - \frac{k_i k_j}{2E}$ to describe the expected number of edges between node pairs, in which $k_i$ and $k_j$ denote the degrees of node $i$ and $j$, respectively. Based on matrix $\mathcal{B}$, the modularity can be calculated as: $Q = \frac{1}{4E}\mathrm{Tr}(\vec{x}^T \cdot \mathcal{B} \cdot \vec{x})$, in which $\vec{x}$ is the column vector describing the partition of a network. Vector $x$ has elements $x_i = \pm 1$ indicating the modules to which the node belongs. For any given partition $s$ of a network, one can calculate $Q$ corresponding to such a partition. The appropriate partition of a network would maximize the modularity $Q$ [22]. In this work, we applied the Louvain algorithm [23] to partition the network and maximize the value of modularity $Q$.

#### 6.3 Dataset

In the file "Protein_Data.txt" (file accessible at: https://bit.ly/37AcczU), the PDB codes for all the proteins in our data set are listed. Besides, the chain length $N$, radius of gyration $R_g$, modularity $Q$, power-law coeffient $\zeta$, spectral dimension $d_s$, total SASA, mean SASA (SASA per residue), $\lambda_1^{-1}$, $\lambda_2^{-1}$, functional sensitivity $1/(\lambda_1\lambda_2)$, mutational robustness $|\lambda_1^{-1} - \lambda_2^{-1}|$, $|\lambda_2^{-1} - \lambda_3^{-1}|$, and $|\lambda_3^{-1} - \lambda_4^{-1}|$. All the calculations are based on $r_C = 8$Å.
